# Desert fish populations tolerate extreme salinity change to overcome hydrological constraints

**DOI:** 10.1101/2021.05.14.444120

**Authors:** Celia Schunter, Lucrezia C. Bonzi, Jessica Norstog, Jade Sourisse, Michael L. Berumen, Yoseline Angel, Stephen D. Parkes, Matthew F. McCabe, Timothy Ravasi

**Author notes:** corresponding authors: Celia Schunter and Timothy Ravasi.

## Abstract

The unstable nature of freshwater ponds in arid landscapes represent a sizable challenge for strictly aquatic organisms, such as fishes. Yet the Arabian Desert, bordering the coastline of the Red Sea, plays host to a species very well adapted to such extreme environments: the Arabian pupfish, *Aphanius dispar*. In this study, we estimated patterns of hydrological connectivity; population structure and stable isotope for samples of *A. dispar* living in small, isolated ponds of nearly-freshwater in the Arabian desert and highly saline coastal lagoons along the Red Sea. The genomic and hydrological analyses indicate that populations are largely separated by drainage origin, as fish from desert ponds appear to be transported to coastal lagoons of the Red Sea along ephemeral river systems arising from flash flood events. Further, our study indicates there is an ecological change when being washed from pond environments to coastal waters, due to a significant shift in muscle stable isotopes ratios between both groups. Considering that the genetic breaks are mostly observed between drainage origin, this study suggests that *A. dispar* can survive large changes in salinity and ecological regimes over small time-scales.

## Introduction

Population dynamics are often defined and restricted by differences in environmental conditions. A growing body of research shows that environmental gradients are often a driver for selection, but also population structure, as adaptation to local conditions can lead to selection of particular genotypes on either side of an ecological break (Wang & Bradburd, 2014). Differences in fitness and conditioning across such ecological gradients can lead to restricted dispersal or even dispersal barriers, which in turn can significantly shape the population structure at relatively small geographic scales (Sexton, Hangartner, & Hoffmann, 2014). This concept is tremendously important in evolutionary biology, as ecological gradients have been suggested as one of the initial mechanisms of divergence that can promote speciation of lineages (Doebeli & Dieckmann, 2003).

A small number of species have the ability to traverse ecological barriers, even significant dispersal barriers and eventually conquer some of the world’s extreme environments, via physiological and behavioural adaptations (Kelley et al., 2014; Moore, Cooper, Biewener, & Vasudevan, 2017). Among the most intriguing of these examples are fishes that live in desert environments, as they inhabit highly isolated and often ephemeral ponds in very arid landscapes. It is assumed that most of these species have high phenotypic plasticity, as these fishes encounter ephemeral streams (Furness, 2016), large temperature fluctuations (Bennett & Beitinger, 1997), extreme changes in water chemistry (Kavembe, Franchini, Irisarri, Machado-Schiaffino, & Meyer, 2015) and high spatiotemporal variability in water supply (Fisher, Gray, Grimm, & Busch, 1982). Among the main questions regarding these particular groups of fishes, the most intriguing is how such isolated populations continue to survive after an initial colonization event. It is well known that isolation of populations can result in a slew of detrimental conditions, such as loss of genetic variability, inbreeding depression or the accumulation of deleterious mutations (Gaggiotti, 2003). Considering that population persistence and long-term survival is largely influenced by genetic diversity (Bouzat, 2010), living in isolated and highly restricted water bodies might threaten the persistence of desert fishes over long time-scales.

Among the few fish groups that inhabit arid environments, Cyprinodontiformes exhibit remarkable adaptations to extreme conditions. For example, the African turquoise killifish (*Nothobranchius furzeri*) evolved an embryonic diapause, allowing fertilized eggs to survive a dry period (Furness, 2016). Furthermore, killifish can also be found in habitats with widely varying salinities and they are therefore categorized as euryhaline (Wood & Marshall, 1994). The genus *Fundulus,* for instance, displays a wide range of osmotolerant physiologies (Whitehead, 2010), with *F. heteroclitus* being able to rapidly acclimate to an osmotic shock by changing its transcriptional program and later remodeling its tissues (Whitehead, Galvez, Zhang, Williams, & Oleksiak, 2011). Despite these remarkable adaptations, some Cyprinodontiformes occupy very restricted habitats. For example, the Devils Hole pupfish (*Cyprinodon diabolis*) lives only in Devils Hole (in the Amargosa Desert of Nevada) and it is described as occupying the smallest known natural range (a single pool < 80 m^2^) for any vertebrate species (Martin, Crawford, Turner, & Simons, 2016). Indeed, many species of fish inhabiting such arid landscapes are currently listed as endangered (Hopken, Douglas, & Douglas, 2013; Van Haverbeke, Stone, Coggins, & Pillow, 2013) and have been of conservation concern for decades (Meffe & Vrijenhoek, 1988).

The Arabian pupfish, *Aphanius dispar* (Cyprinodontidae), is present in the Middle East with landlocked populations in countries such as Oman, Iran and Saudi Arabia (Al-Kahem-Al-Balawi et al., 2008; Freyhof, 2014; Haas, 1982; Hrbek & Meyer, 2003). This species represents an interesting conundrum, as it has a large distribution range, but it inhabits highly restricted ephemeral ponds with no permanent rivers around them. Furthermore, from a pilot study we revealed that they are also present in coastal lagoons of the Red Sea. Phylogenetic analyses of the genus suggest that this group has saltwater ancestry, with the closing and drying of the Tethys Sea resulting in landlocked remnant populations which likely then diverged due to the resultant strong ecological changes (Hrbek & Meyer, 2003).

In this study we explore suitable habitats for *A.dispar* to live in the desert as well as the Red Sea coastline in Saudi Arabia and aim to understand the population connectivity of this species. We investigate the environmental conditions and population structure of *A. dispar* in the Arabian desert by employing a multi-disciplinary approach that combines hydrological predictive mapping, population genomics, stable isotope analysis, as well as chemical and physical analyses of water bodies.

## Methods

### Sample collection

In order to acquire information on the presence of water ponds and streams in the Saudi Arabian desert, extensive searches were performed using Google Earth, high-resolution satellite data and through contacting local landowners and regional police. Subsequently, a range of potential water ponds were identified and visited (see Figure 1). From a survey of 28 locations, *Aphanius dispar* was detected at 11 sites and was collected using a 3 m wide seine net. Fish length was measured, and a piece of the dorsal fin was cut and placed in 96% ethanol. Fish were then returned to the pond. Individuals with fins too small for finclip collection were euthanized with a blow to the head, with the entire tail preserved in ethanol. All procedures were performed in accordance with relevant guidelines and regulations and were approved and completed with the ethics permit 15IBEC35_Ravasi from the Institutional Biosafety and BioEthics Committee (IBEC) of the King Abdullah University of Science and Technology.

**Figure 1:**
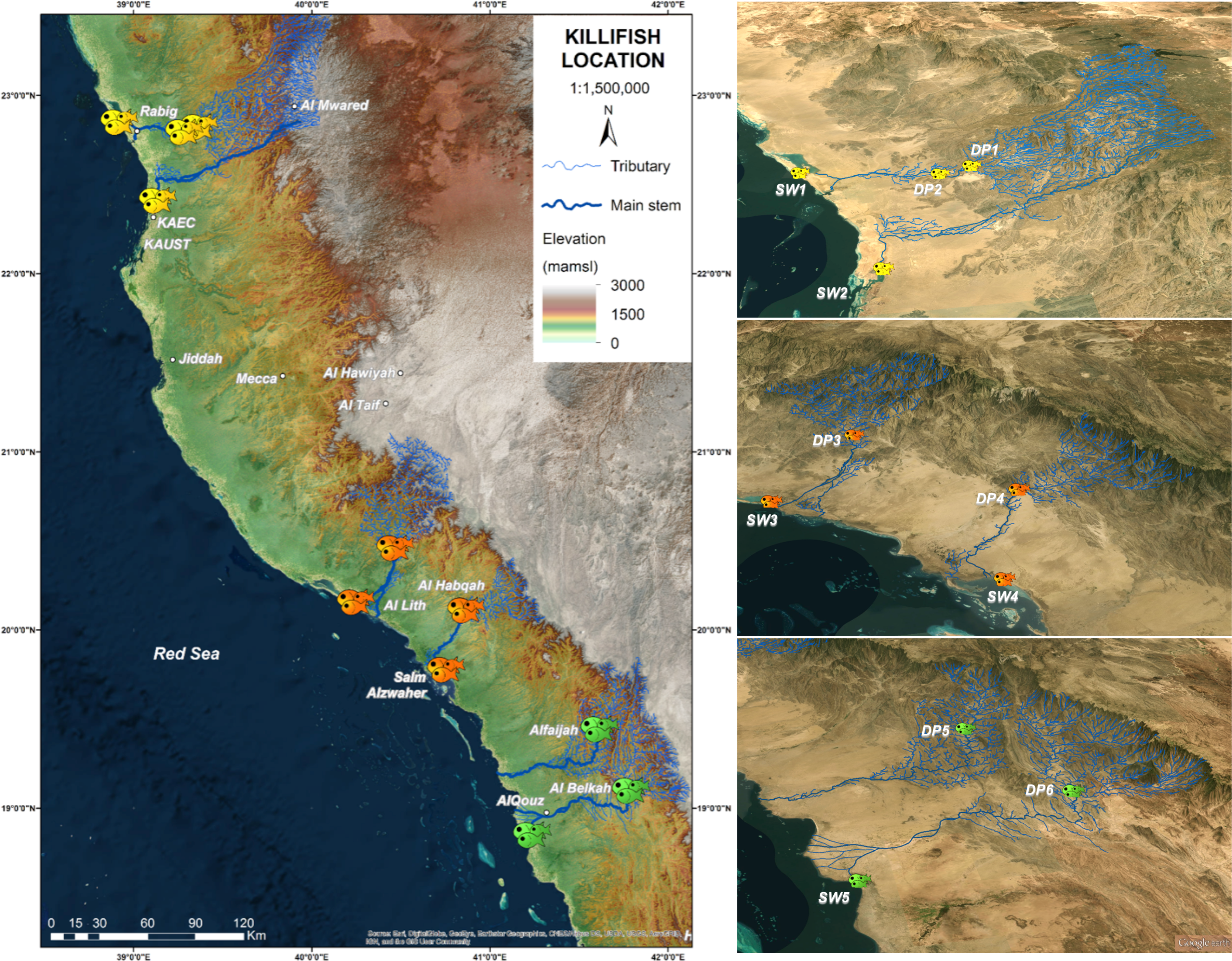
Locations of all sampling sites over a 1,100 km stretch of the Saudi Arabian coastline. Grey circles represent desert ponds without the presence of *Aphanius dispar*. Sites with pupfish are colored (n=11) and classified as desert ponds (DP) or seawater lagoons (SW). The top two pictures show a male and female *A. dispar*, respectively. The center picture is an example of a desert pond (DP2), below which is the Al Lith hotspring (DP3). The bottom picture shows an example of a seawater site (SW4). The map was created using ArcGIS 10.5 (www.arcgis.com).

### Stable isotopes

We employed two types of stable isotope measurements: fish muscle tissue and pond water isotopes. Fish muscle tissue can give indications on the diet of the fish in the ponds and the sea (Zanden & Rasmussen, 2001). Water isotope analysis can indicate the evaporative and flow processes of the water body. While these distinct isotopic measurements have different applications, they both help to understand the biological and hydrological connectivity of the ponds in the desert. Isotope ratios are reported relative to their respective standards:

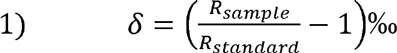

where R_sample_ and R_standard_ are the isotope ratios (heavier isotopologue to the more abundant isotopologue) measured in the sample.

#### a) Water stable isotope analysis

Water samples from the six desert ponds were analyzed for isotope analysis at the King Abdullah University of Science and Technology on a Picarro Wavelength Scanning Cavity Ringdown Spectrometer (WSCRDS L115-I, Picarro Inc., Sunnyvale, CA, USA) interfaced to a liquid autosampler (CTC HTC Pal liquid autosampler; LEAP Technologies, Carrboro, NC, USA). δ^2^H and δ^18^O were measured and are isotope ratios for ^2^H^1^H^16^O and ^1^H_2_^18^O isotopologues, respectively. Samples were referenced to the VSMOW scale using laboratory working standards previously referenced to the VSMOW2/ SLAP2 (Standard Light Antarctic Precipitation) standards. The δ^2^H and δ^18^O values of samples was determined using the two-point linear normalization method described in Paul *et al*. (2007) (Paul et al., 2007), which calibrate sample isotope ratios using the linear relationship between the true and measured isotope ratios of two working standards that bracket the sample. Each measurement of standards and samples consisted of 10 injections of which the last 4 were used to determine isotope ratios. Uncertainty for the reported isotope values is determined by propagating the standard deviations (1σ the last 4 injections from measurement of standards and samples.

#### b) Muscle tissue stable isotope preparation and analysis

For each collection site, a subset of adult fish was sacrificed, and bodies were frozen with dry ice in the field. In the lab, fish were thawed, and white muscle tissue was de-skinned, descaled and dried for 24 h at 60 °C. Samples were ground using a MP Bio FastPrep-24 instrument (6.5 m/s for 60s until pulverized, repeated 1-4 times) and Lysing Matrix E 2 mL tubes with 1 x 4.0 mm ceramic sphere, 30±3 1.4 mm ceramic spheres. Tissues were rinsed in 1 mL 2:1 chloroform-methanol solution and shaken vigorously for 30 s to remove isotope differences due to a lipid content bias, as described by Ehrich *et al*. (2011) (Ehrich et al., 2011). The solution was transferred to 1.5 mL Eppendorf tubes, leaving the spheres behind. The samples were left to stand for 15 minutes at room temperature, and then centrifuged for 10 minutes at 3,400 rpm. The supernatant was discarded, and the chloroform-methanol rinse was repeated two additional times. Samples were then dried in 1.5 mL Eppendorf tubes for 24 h at 60 °C and broken apart using a scoopula. Approximately 1±0.2 mg of sample was placed in tin capsules for solid samples (5 x 9 mm) for the Isotope Ratio Mass Spectrometry analysis (IRMS). ^13^C and ^15^N isotopes were analyzed using a PDZ Europa ANCA-GSL elemental analyzer interfaced to a PDZ Europa 20-20 isotope ratio mass spectrometer (Sercon Ltd., Cheshire, UK) at the Stable Isotope Facility at the University of California, Davis. Briefly, samples were combusted at 1000°C with chromium oxide and silvered copper oxide, with a subsequent oxides removal in a reduction reactor. N_2_ and CO_2_ were separated on a Carbosieve GC column (65°C, 65 mL/min) before entering the IRMS. Samples were interspersed with replicates of two laboratory standards and isotope ratio delta values were measured relative to the international standards VPDB (Vienna PeeDee Belemnite) and Air for carbon and nitrogen, respectively (Sharp, 2007). Statistical analyses for tissue stable isotopes were carried out using R (R version 3.4.0, 2017-04-21). Data were checked for normality with the Shapiro-Wilk test and for homogenous variance with the Fligner-Killeen test and considered significant at *p* < 0.05. The correlation between δ^13^C and δ^15^N was tested with a Spearman’s correlation, as well as the allometric effect of size on isotope contents. Since data did not satisfy conditions for the use of parametric statistics, a two-way non-parametric ANCOVA in the *sm* package (Bowman & Azzalini, 2014) was used to test for differences in δ^13^C and δ^15^N among sites, with fish size as the covariate. Sampling sites of *A. dispar* were split into two water type groups (Desert Pond and Sea Water), as well as in four different clusters (determined through genetic and hydrological analysis, stated below), depending on the drainage basin they belong to (from north to south: Cluster 1 = DP1, DP2, SW1, SW2; Cluster 2 = DP3, SW3; Cluster 3 = DP4, SW4; Cluster 4 = DP5, DP6, SW5). Differences in fish muscle isotopic signatures between different desert pond were also evaluated. Muscle mean content in δ^13^C and δ^15^N and standard deviation for the different sites (n: DP3 =11, DP4 = 11, all others = 12) were calculated and visualized in R.

### Water parameters and samples

To quantify the water chemistry of the desert ponds and better understand the environmental conditions of the fish habitat, we took *in situ* measurements as well as collected water samples for later analyses. A portable Ocean Seven 305 Plus CTD (Idronaut) was used to measure temperature, pressure, conductivity, salinity, percent oxygen saturation, parts per million of O_2_ and pH on site. The CTD sonde was submersed at a depth of approximately 30 cm and left to acclimate for one minute. Five minutes of measurements were taken at a sampling rate of 5 seconds, with the total of 60 data points averaged to provide a single value for each parameter per site (referred to as “average temperature”). Water samples at each site were collected in 500 ml HDPC containers, either filled only with water for isotope and water analysis, or with 2ml of 5% nitric acid for preservation for chemical water analysis. Water samples were analyzed using Ion Chromotography. Seawater samples (sites SW1 – SW5) were diluted x10 and filtered using Dionex OnGuard II Ba/Ag/H 2.5 cc cartridges to remove SO_4_^2-^ and Cl^-^. Standards and water samples were transferred to Autoselect Polyvial 10 mL vials and covered with septa and caps. Samples were run on Dionex ICS-3000 with the Chromeleon Chromatography Management System (version 6.7) program and using an autosampler. Photometric analysis was performed on water samples to measure for chlorides, sulphates and silica. Standards were run accordingly (chlorides: 0 ppm, 20 ppm, 200 ppm; sulphates: 0, 30 and 100 ppm; silica: 0, 100 and 500 ppb). Water samples were transferred to cuvettes and loaded into an Aquakem 250 (Thermo Scientific) for analysis. For samples that were outside of standard ranges, samples were diluted appropriately (see Supplementary materials) and re-run.

### Streamflow mapping

To establish the hydrological connectivity from the desert highlands to the Red Sea coastline (and the resultant possible genetic links between the locations where pupfish were present), we needed to determine the predominant flow direction at each cell of an underlying topographic model. The ArcGIS-based hydrology toolset was used for the extraction and analysis of watersheds and streamflow, using the Advanced Spaceborne Thermal Emission and Reflection Radiometer (ASTER) Global Digital Elevation Model (ASTGTM, JPL 2009) to provide a topographic description of the region. Raw ASTER data, distributed by NASA as GeoTIFF files, have a 30 m spatial resolution and are referenced to the WGS84 coordinate system. Supplementary Figure 1 shows the flow process of delineating watershed boundaries and stream networks from a digital elevation model (DEM). To ensure that any water “flow” can move from one cell to any adjacent cell, a depressionless DEM was obtained by filling any localized “sinks” that might have formed as an artefact of the interpolation process. From this, the natural flow direction (as dictated by the direction of steepest descent) and the flow accumulation per cell (i.e. the number of upstream cells “draining” to that particular cell) can then be calculated. Only those cells with a high accumulation threshold (>1000 contributing cells) are considered to represent a dominant flow path. A streamline can be produced by connecting these cells. Watershed boundaries are delineated automatically based on the natural water divides, which follow the highest elevations in the DEM. Using Google Earth imagery (Google Earth 7.1.2.2041; December 31, 2016) as an underlying base map, we can determine the potential connectivity routes for fish by following delineated streamlines from the highest sampled locations on the mountains, to the lowest points near the shoreline. A visual interrogation of satellite images allowed elements such as dams, bridges, culverts and agricultural regions to be identified and to manually edit segments along the streamlines.

### DNA extractions and Restriction Site Associated DNA Sequencing (RAD-Seq)

To understand the genetic population structure and genetic connectivity of *A. dispar* we used 5 mm pieces of fin clip from each individual sample for 28-29 individuals per site (at 11 sites) to extract DNA with 96-well DNA extraction kits from Qiagen (DNeasy 96 Blood and Tissue Kit) or Macherey-Nagel (NucleoSpin 96 Tissue). The manufacturer protocols were followed with a deviation of 8-10 hours of lysis and elution in 50 μl H_2_0. Concentrations were measured with a Qubit 2.0 fluorometer with a dsDNA High Sensitivity reagent kit. We used a modified double digest (ddRad) protocol (Peterson, Weber, Kay, Fisher, & Hoekstra, 2012). Briefly, DNA was digested using SphI High Fidelity and MluCI High Fidelity enzymes with the appropriate CutSmart Buffer (New England Biolabs) and cleaned with AMPURE XP beads. Adapters were ligated to 100 ng of digested DNA with a combination of sixteen adapters and eleven indices to uniquely identify individual samples out of 80 multiplexed samples in one sequencing lane. Pools were created from equally concentrated and cleaned samples and size selected in a BluePippin (Sage Science) with 2% Gel Cassettes to a size of 300bp. We used a KAPA Hifi Ready Mix for the PCR amplification with ten PCR cycles and temperatures according to the manufacturer. Samples were run on a bioanalyzer (Agilent) and a qPCR (7900 HT Fast Real Time PCR system, ABI) was used to quantify and combine all 80 samples within one library at equimolar concentration. Four libraries were then sequenced paired-end on an Illumina Hiseq2000 to a length of 100bp at the KAUST Bioscience Core Lab facility.

### RAD-Seq data processing

Raw sequence fastq files were de-multiplexed for each lane of Illumina using *process_radtags* in the software STACKS 1.40 (Catchen, Hohenlohe, Bassham, Amores, & Cresko, 2013). All sequences were quality trimmed with Trimmomatic 0.33 (Bolger, Lohse, & Usadel, 2014) with a Phred score quality cutoff of 30. Individuals with less than 300,000 remaining first read sequences (from paired end reads) were removed from the analysis. Due to the lack of a reference genome for this species, we ran the quality trimmed first read files through the *denovo_map* perl script in STACKS. The final optimized parameters were set to a conservative number of mismatches allowed (-n 2 and -M 2), a minimum number of identical reads to form a stack was three and SNP calling was performed with an upper bound error rate of 0.05 (--bound_high 0.05). After the putative SNP detection, the function *populations* was run to select putative SNPs meeting several criteria. The variant had to have a minimum read number of 10 (-m 10), be present in at least seven of the eleven locations (-p 7) and in more than 20 individuals per locations (-r 0.72). A minimum allele frequency filter of 0.05 was applied and we chose to use one randomly putative SNP from each stack. The resulting vcf file was converted into different input file formats for further analysis using PGDSPIDER (Lischer & Excoffier, 2012). To avoid a bias due to duplicated sequences across the genome, putative SNPs were discarded if heterozygosity was higher than 0.5 (calculated with *vcftools,* following (Danecek et al., 2011) and if the mean read depth was a median absolute deviation away from the median depth (Seeb et al., 2014). Hardy-Weinberg exact tests were performed in Genepop version 4.6 (Rousset, 2008) and putative SNPs were removed if there was a significant deviation in more than four of the eleven locations for which tests could be performed. All SNPs that did not pass the criteria were blacklisted in *populations* and removed from further analysis.

### Population genetics and clustering analyses

Population genetics metrics such as average allele number, private alleles, inbreeding coefficient and locus and pair-wise F_ST_ were obtained for the set of filtered final SNPs with *populations*. VCFtools v0.1.13 was used to look at minor allele frequencies, Hardy-Weinberg equilibrium and locus and population heterozygosities (Danecek et al., 2011). Pairwise AMOVA F_ST_ values for population differentiation measures and significance (p<0.05) were obtained in GenoDive V2.0b27 using 10,000 permutations as well as the pairwise kinship coefficient r (Meirmans & Van Tienderen, 2004). Effective population sizes were calculated in NeEstimator using the linkage disequilibrium method including all final selected SNP loci (Do et al., 2014). Here we present the results of the largest harmonic mean sample size per population.

To evaluate genetic structure among all sampled individuals, we performed a principal component analysis (PCA) as well as Bayesian clustering analysis. The PCA was computed in the *adegenet* package v. 2.0.1 in R (Jombart, 2008). We represent the eigenvalues of the analysis revealing the variance of each principal component and a scatterplot summarizing the genetic diversity including the center ellipses per sampling location. Clustering analysis was performed in Structure v.2.3.4 (Pritchard, Stephens, & Donnelly, 2000) under the admixture model with a 10% burn-in period and 500,000 iterations of Markov Chain Monte Carlo (MCMC), which creates a probability distribution and allows for the evaluation of the likeliness of different numbers of clusters (K) within the dataset. Ten replicates were run for each putative number of clusters (K) with K ranging from 1 to 11. The results were then passed through STRUCTURE HARVESTER v0.6.94 (Earl & vonHoldt, 2012) to apply the *ad hoc* statistic delta K proposed by Evanno and coauthors (2005). Resulting individual and population files were used in CLUMPP v1.1 (Jakobsson & Rosenberg, 2007) and DISTRUCT v1.1 (Rosenberg, 2003) to combine all STRUCTURE runs and visualize the results.

### Loci under selection

As *A. dispar* individuals were found in very different habitat types (from desert ponds to the highly saline waters of the Red Sea) we tested for possible selection in any of the analyzed loci possibly indicating adaptive processes to the environmental conditions. For this we re-ran the *populations* program with the same whitelist of 5,955 loci selecting specific locations. First, all locations were used by defining all desert pond locations as one population and all seawater location as another one, in order to evaluate global selection between desert and seawater habitats. However, adaptive processes might differ for the different desert ponds and we therefore sub-selected locations. We evaluate adaptive loci for DP1 & DP2 against SW1 & SW2, DP3 vs. SW3, DP4 vs. SW4, and DP5 & DP6 against SW5. Each resulting vcf files was format converted with PGDSpider 2.0.0.2 (Lischer & Excoffier, 2012). For outlier loci detection we used a Bayesian approach incorporated in Bayescan v2.1 (Foll & Gaggiotti, 2008). Briefly, posterior odds of a locus being under selection are obtained with MCMC with the help of the proportion of loci exhibiting large F_ST_ in comparison to other loci. To further minimize the number of false positives, the prior odds was increased to 100 (-pr_odds 100). Outlier loci were then visualized and selected in R after applying a False Discovery Rate (FDR) of 0.05. Sequence reads for resulting putative loci under selection were then blasted against the NCBI nr database (blastn) and successful blast hits are presented if the evalue was below 1e^-5^.

## Results

### Pupfish locations, collection and pond properties

Due to the lack of previous knowledge on the locations of the Arabian pupfish populations in western Saudi Arabia, we sampled 28 sites, of which only eleven had pupfish present (Figure 1). 12 Red Sea coastline sites and 16 inland enclosed ponds or streams were visited (Table 1 & Supplementary Table 1); 6 inland sites and 5 seawater sites had *Aphanius dispar* specimen. For two inland ponds, no sonde measurements could be obtained due to complications transporting the CTD (conductivity, temperature and depth) sonde into the steep canyons. Five ponds contained freshwater with a salinity upper maximum threshold of 0.5 ppt, whereas salinity in the other nine measured ponds ranged from 0.54 to 1.45 ppt. Most of the ponds therefore consisted of brackish water. *Aphanius dispar* was only found in ponds with salinities higher than 0.74 ppt. pH ranged between 7.71 to 9.41 across all sites, with the majority exhibiting a pH between 8 and 9. The average temperature in the inland ponds was 28.9 °C (±4.9 SE), while in the Red Sea sites it was 21.9 °C (±4.17 SE). Sampling of fish was performed during the late boreal autumn and early winter months (i.e., November-January), hence during the period of lowest temperature of the year. The highest average water temperature, 38.4 °C, was observed at the Al Lith hot spring. At this site, pupfish live in higher temperatures in comparison to all other sites, as well as with larger water content of silica (80,470 ppb). These results illustrate the wide range of environments which pupfish populations are able to exploit along the Red Sea coast.

**Table 1:** Sampling locations, characteristics and selected water parameters (for full water measurements refer to Supplementary Table 1) measured by CTD or Aquakem. A dash indicates that no reliable measurement was obtained.

| Site | Type | Temperature (°C) | Conductivity (µS/cm) | Salinity (ppt) | O2 (ppm) | pH | Chlorides (ppm) | Silica (ppb) | pupfish present |
| --- | --- | --- | --- | --- | --- | --- | --- | --- | --- |
| SW5 | Sea | 21.77 | 58,340.47 | 41.93 | 2.87 | 8.71 | 27.10 | 306.02 | yes |
| SW4 | Sea | 27.90 | 66,952.99 | 42.75 | 3.39 | 9.01 | 29.59 | 314.61 | yes |
| SW3 | Sea | 22.82 | 58,685.99 | 41.19 | 4.40 | 9.10 | 2.15 | 104.57 | yes |
| SW2 | Sea | 24.80 | 64,966.91 | 44.20 | 3.43 | 8.37 | 36,677.27 | 517.96 | yes |
| SW1 | Sea | 20.90 | 59,185.03 | 43.49 | 3.93 | 8.37 | 36.06 | 31.97 | yes |
| DP6 | Pond | 31.42 | 1,663.76 | 0.74 | 7.75 | 9.41 | 282.41 | 34,511.49 | yes |
| DP5 | Pond | 30.93 | 1,760.39 | 0.79 | 8.32 | 9.39 | 294.11 | 59,944.24 | yes |
| DP4 | Pond | 26.44 | 2,004.49 | 0.99 | 6.14 | 8.31 | 278.57 | 63,358.87 | yes |
| DP3 | Hot Spring | 38.42 | 1,916.49 | 0.75 | 5.16 | 8.20 | 361.21 | 80,469.67 | yes |
| DP2 | Pond | 31.58 | 3,190.00 | 1.45 | 4.35 | 8.31 | 734.37 | 38,017.18 | yes |
| DP1 | Pond | 29.93 | 1,720.00 | 0.78 | 4.35 | 8.50 | 469.77 | 33,586.98 | yes |
| 28 | Sea | 27.65 | 64,108.41 | 40.93 | 3.32 | 8.46 | - | - | no |
| 27 | Sea | 21.77 | 65,110.01 | 47.50 | 3.81 | 8.21 | - | - | no |
| 26 | Sea | 18.18 | 51,228.77 | 39.44 | 4.56 | 8.40 | - | - | no |
| 25 | Sea | 22.60 | 57,621.28 | 40.54 | 3.61 | 8.48 | - | - | no |
| 24 | Sea | 12.08 | 46,285.24 | 41.12 | 4.11 | 8.29 | - | - | no |
| 23 | Sea | 21.74 | 63,528.52 | 46.35 | 3.86 | 8.24 | - | - | no |
| 15 | Pond | 21.28 | 5,825.19 | 0.31 | 6.00 | 8.60 | 76.24 | 20,016.72 | no |
| 14 | Pond | 21.15 | 8,023.78 | 0.43 | 3.10 | 7.71 | 115.34 | 38,414.86 | no |
| 13 | Pond | - | - | - | - | - | 103.07 | 5,016.26 | no |
| 11 | Pond | 28.91 | 898.74 | 0.41 | 6.33 | 8.04 | 86.54 | 38,664.80 | no |
| 9 | Pond | 33.54 | 1,025.76 | 0.43 | 6.26 | 9.15 | 127.94 | 52,096.62 | no |
| 7 | Pond | 30.21 | 1,257.23 | 0.56 | 4.49 | 8.05 | 196.26 | 76,975.60 | no |
| 6 | Pond | 28.31 | 1,164.95 | 0.54 | 4.69 | 8.34 | 128.02 | 46,885.01 | no |
| 5 | Pond | 30.25 | 879.72 | 0.39 | 4.26 | 8.11 | 135.60 | 25,711.03 | no |
| 4 | Pond | - | - | - | - | - | 29.70 | 10,464.11 | no |
| 3 | Pond | 21.60 | 1,019.11 | 0.54 | 5.32 | 8.25 | 166.66 | 45,331.00 | no |

### Water stable isotopes

To further understand environmental conditions of the six inland ponds inhabited by *A. dispar,* water samples were collected, and their stable oxygen and hydrogen isotopes measured (Supplementary Table 1). This can aid in understanding the source of the water in the desert pond and if evaporation occurs comparatively, which can indicate water flow or standing waters. A strong linear relationship between the δ^2^H and δ^18^O of these samples was observed, with a slope of 4.83. Supplementary Figure 2 shows the linear regression of measurements, with data exhibiting a low slope compared to the Global Meteoric Water Line (GMWL). The relatively low slope indicates samples were subject to significant evaporative enrichment (Gat, 1996; Gibson, Birks, & Edwards, 2008; Gibson & Reid, 2014). The linear relationship between δ^2^H and δ^18^O across all samples indicates a common water source derived from a single recharge event, with variability between samples caused by the degree of evaporative enrichment for each water pool. This is consistent with observations made at each site, with DP1 and DP2 being the two most stagnant ponds with the least oxygen and greatest δ^2^H and δ^18^O isotopic enrichment (Table 1).

### Streamflow mapping

The overall topography of the study region is a mountainous inland region that is bounded to the east by the Arabian shield, and which drains westward to a flatter desert terrain towards the Red Sea coastline. In order to understand the hydrology of the area and the exact hydrological constraints for the Arabian pupfish we created streamflow maps for each sample location derived from a satellite-based digital elevation model. Using the derived stream networks, a total of 6 different migration pathways were identified within the study area (Figure 2 & Supplementary Table 4). Figure 2 illustrates the digital elevation model (DEM) based hydrological connectivity as streamlines flowing from each of the desert site locations where pupfish were present. We find that during periods of occasionally intense or sustained rainfall, the desert ponds may become hydrologically connected to their downstream saltwater location through tributary and main stem water flow. Four distinct areas of potential hydrological connectivity can be determined from Figure 2. In the northern portion of our study region, water from DP1 and DP2 are hydrologically linked via defined streamlines to SW1. SW2 serves as a regional seawater sample pair for SW1, as it is separated by a distance of approximately 40 km. Further south, there is delineated connectivity between DP3 and SW3, as well as between DP4 and SW4. In the southernmost part of the sampling area, DP6 has a tributary stream that connects with SW5, while DP5 has a separate watercourse to the sea. It is important to note that these streamlines do not represent active flow paths, as Saudi Arabia has no permanent rivers. Instead, the streamlines describe either permanently or intermittently dry riverbeds, defined as a function of the topography, with ephemeral flows only occurring in cases of sufficient rainfall.

**Figure 2:**
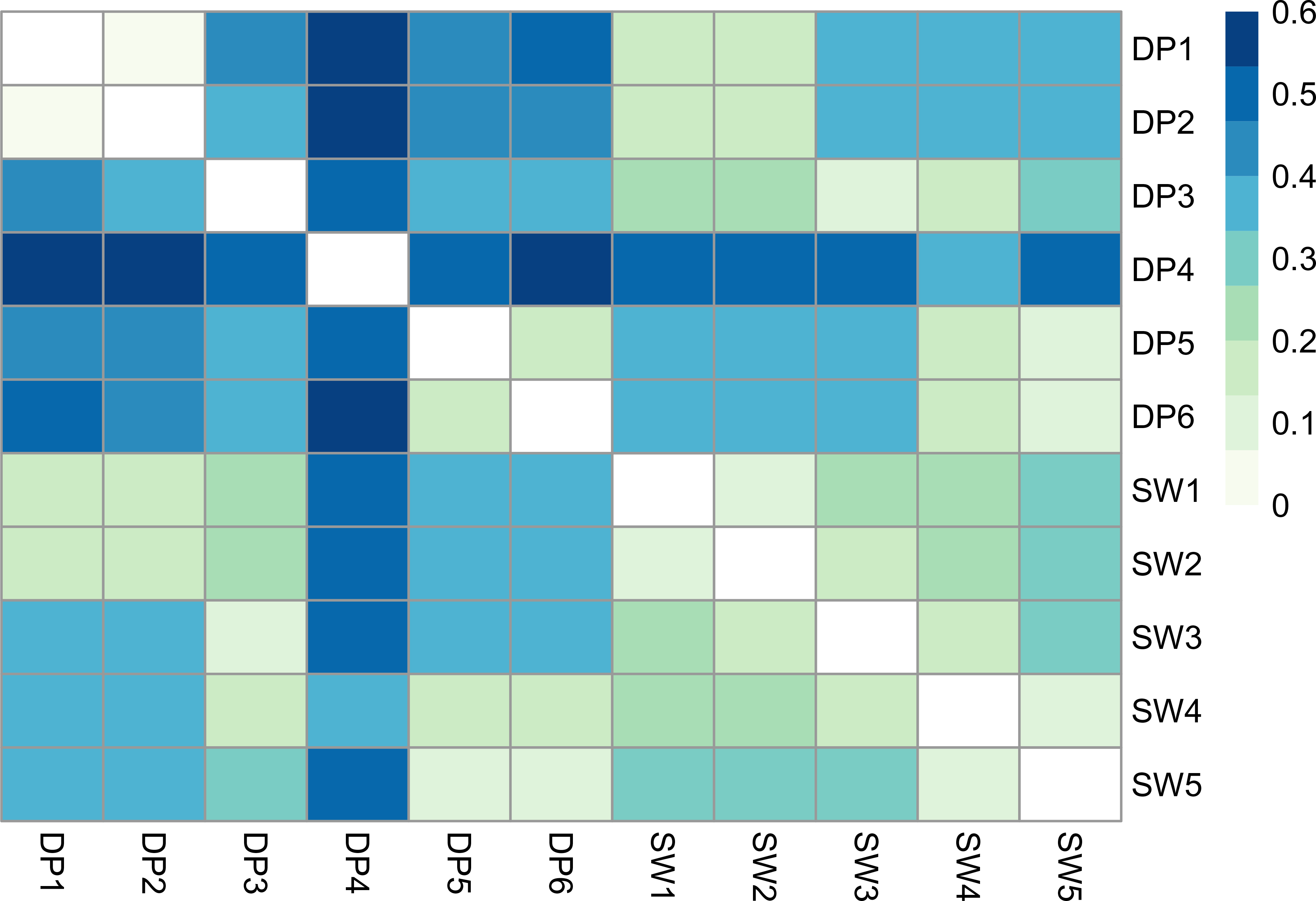
Hydrological modeling. Latitudinal 2D overview of Saudi Arabian coastline with *Aphanius dispar* collection sites indicated with fish icons (left) and 3D close-ups on three regions with predicted hydrological flow. Maps were produced with Google Earth imagery (Google Earth 7.1.2.2041; December 31, 2016) and ArcGIS 10.5 software (www.arcgis.com).

### RAD-sequencing

In order to verify the potential hydrological connectivity for the Arabian pupfish, we use Single Nucleotide Polymorphism (SNPs) genetic markers to evaluate the genetic population connectivity of *Aphanius dispar*. For this we genotyped between 28 and 30 individuals per site for the eleven *A. dispar* locations by means of RAD-sequencing (Table 2 & Supplementary Table 2). Over 2 million sequence reads were obtained on average for each individual. Six samples were removed from further analysis as they had fewer than 300,000 reads after demultiplexing and quality trimming, resulting in 27 to 30 individual samples per location. The total final sample number was 314 individuals for 11 sites (five saltwater and six desert pond localities) (Figure 1). A total of 690,084 putative single nucleotide polymorphisms (SNPs) were obtained with STACKS (Catchen et al., 2013), which were stringently filtered to a final 5,955 SNP loci. The average depth of coverage for these loci was 48x, with a minimum of 24x and an average minor allele frequency of 0.21 (Supplementary Figures 4 & 5).

**Figure 3:**
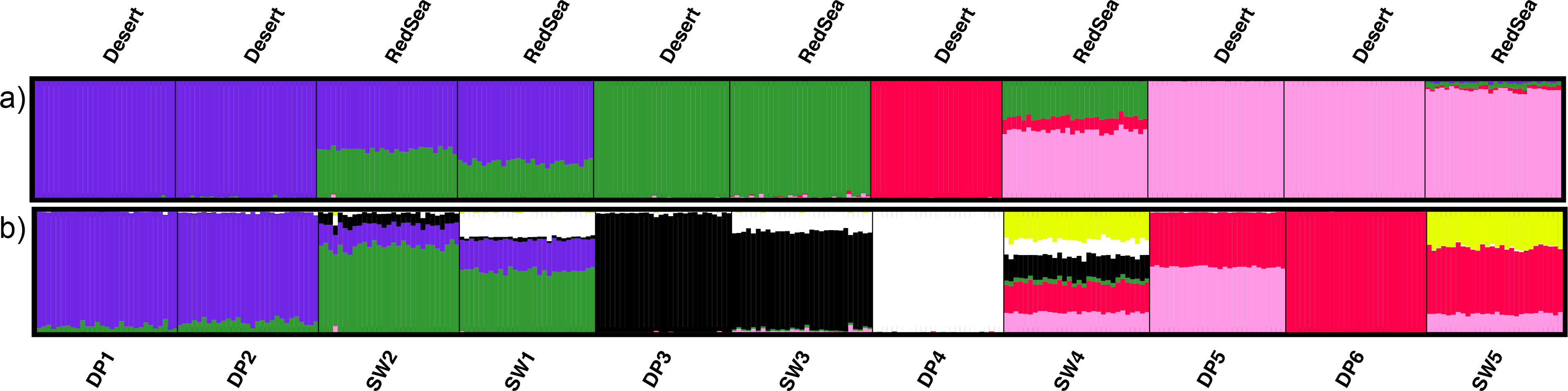
Heatmap of F_ST_ values of pairwise population comparisons based on 5,955 SNPs. Location abbreviations as in Table 1 (DP=desert pond, SW=seawater).

**Figure 4:**
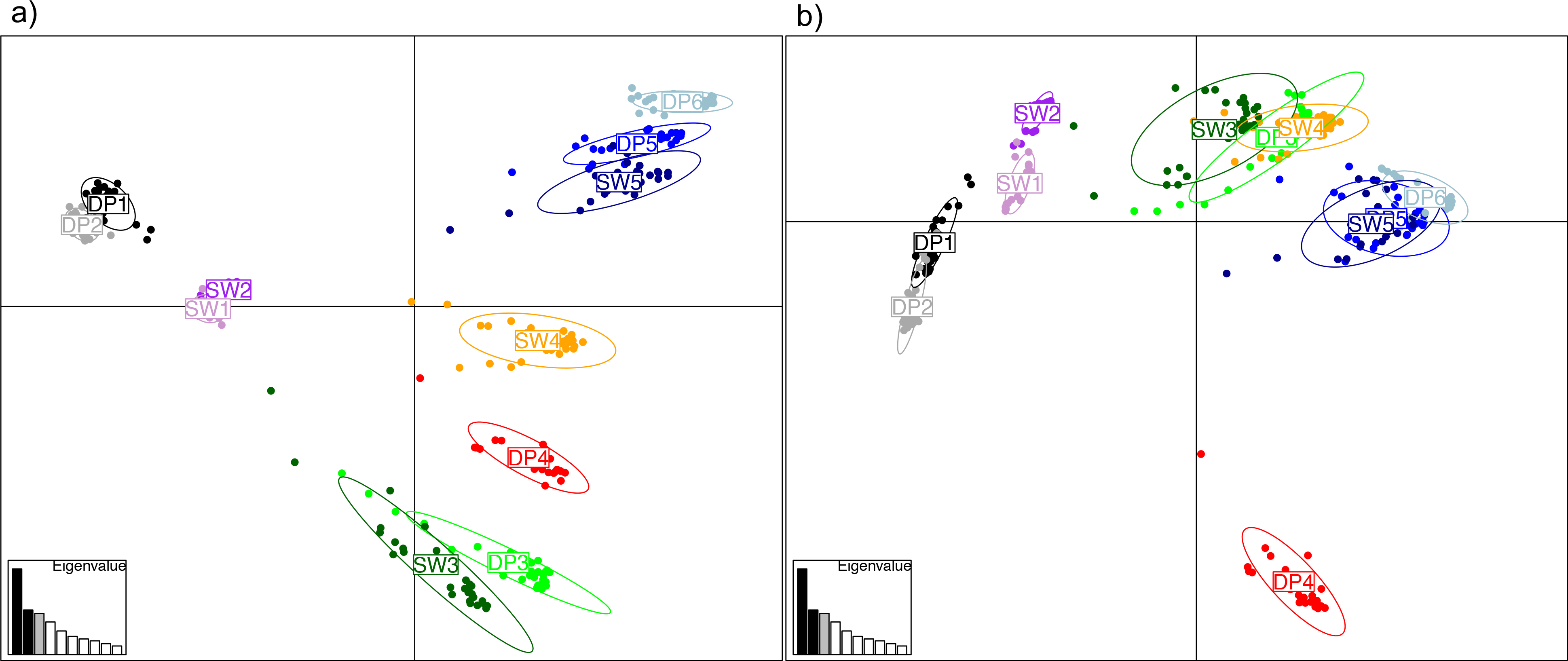
*Aphanius dispar* population structure with a) K=4 and b) K=8. Each individual in the different sites (y-Axis) are plotted with colours representing the probability of each individual to be assigned to a certain cluster.

**Table 2:** Genetic metrics for all 11 pupfish locations for 5,955 SNP loci per sampling site. N = number of individuals, Private a = number of private alleles, Na = average number of alleles, Ho = observed heterozygosity, He = expected heterozygosity, F_IS_ = inbreeding coefficient, Ne = effective population size, 95% CI = 95% confidence intervals for Ne, mean r = average of pairwise kinship coefficient.

| Location | N | Private a | Na | Ho | He | $F_{IS}$ | Ne | 95% CI | Mean r |
| --- | --- | --- | --- | --- | --- | --- | --- | --- | --- |
| DP1 | 29 | 15 | 1.74 | 0.21 | 0.23 | 0.07 | 1222 | 983-1614 | 0.22 |
| DP2 | 29 | 8 | 1.78 | 0.24 | 0.25 | 0.05 | 1851 | 1256-3511 | 0.12 |
| DP3 | 28 | 6 | 1.57 | 0.17 | 0.19 | 0.07 | 464 | 421-515 | 0.12 |
| DP4 | 27 | 27 | 1.31 | 0.08 | 0.08 | -0.02 | 51 | 49-52 | 0.20 |
| DP5 | 28 | 4 | 1.53 | 0.16 | 0.17 | 0.06 | 1497 | 1084-2416 | 0.25 |
| DP6 | 29 | 9 | 1.50 | 0.15 | 0.16 | 0.09 | 1713 | 1236-2782 | 0.13 |
| SW1 | 28 | 7 | 1.80 | 0.24 | 0.27 | 0.08 | 1149 | 882-1646 | 0.27 |
| SW2 | 29 | 13 | 1.80 | 0.23 | 0.26 | 0.09 | 5910 | 2229-inf | 0.06 |
| SW3 | 29 | 0 | 1.66 | 0.19 | 0.20 | 0.06 | 1702 | 1199-2925 | 0.15 |
| SW4 | 30 | 0 | 1.77 | 0.20 | 0.22 | 0.10 | 1645 | 1284-2286 | 0.19 |
| SW5 | 28 | 0 | 1.70 | 0.18 | 0.20 | 0.12 | 3917 | 2175-19493 | 0.42 |

### Population genomics

The number of alleles across the 5,955 SNP loci ranged from 1.3 to 1.8 for the different locations (Table 2). Heterozygosity revealed lower levels for desert ponds and particularly for desert pond DP4 the lowest heterozygosity among all locations, while also having a negative inbreeding coefficient (F_IS_). DP4 also exhibited the most private alleles (27) followed by DP1 with 15 and SW2 with 13. Three seawater locations SW3-SW5 had no private alleles. Pairwise genetic distance between populations (F_ST_) was highest for DP4, with almost all values above 0.2 and the closest genetic distance with SW4 (Figure 3). The lowest value (0) was observed for DP1 and DP2. Interestingly, most DP locations showed higher F_ST_ values when compared to other DP localities than with SW locations, indicating a clear disconnection between desert pond sites. Exceptions to this include DP5 and DP6, with an F_ST_ value of 0.048. Pairwise F_ST_ comparisons between seawater localities ranged between 0.005 to a maximum of 0.105 (SW2 vs SW5). Effective population size ranged between a very low 51 in DP4 to nearly 6,000 in SW2 (Table 2). DP3 was the only other site with an effective size below 1,000.

Bayesian clustering and *ad hoc* testing allowed for the approximation of the most likely number of clusters within the analyzed samples. The most probable cluster numbers were 2 or 8, but delta K *ad hoc* testing also showed a peak at K=4 (Supplementary Figure 6). K=2 divides the northern localities (DP1, DP1& SW1, SW2) from the southern sites, with DP3 and SW3 being an admixed group. When considering four different genetic units in our data set, a division from north to south is found, combining desert ponds and seawater locations together into clusters (Figure 4). In this scenario, DP4 stands out as the only locality with its own cluster. SW4 in turn seems to have the most genetically admixed individuals of all locations. By increasing the cluster number to 8, a further subdivision in the northern part can be found, separating the two desert ponds from the seawater locations (DP1 & DP2 and SW1 and SW2). Here SW4 and SW5, the two furthest southern seawater sites, appear to share some unique traits (represented by the yellow cluster in Figure 4b), which are not shared with the southern desert ponds.

### Hydrological and genetic connectivity

The population clustering in the principal component analysis revealed a break between the four northern locations (SW1, SW2, DP1 and DP2) and the seven southern ones, regardless of them being desert ponds or seawater sites (Figure 5a). The two desert ponds DP1 and DP2 closely clustered together as well as the two seawater locations (SW1 and SW2), with some distance between the two different environments. This disjunction, however, is not seen through hydrological mapping, as DP1 & DP2 have hydrological connectivity to SW1.

**Figure 5:**
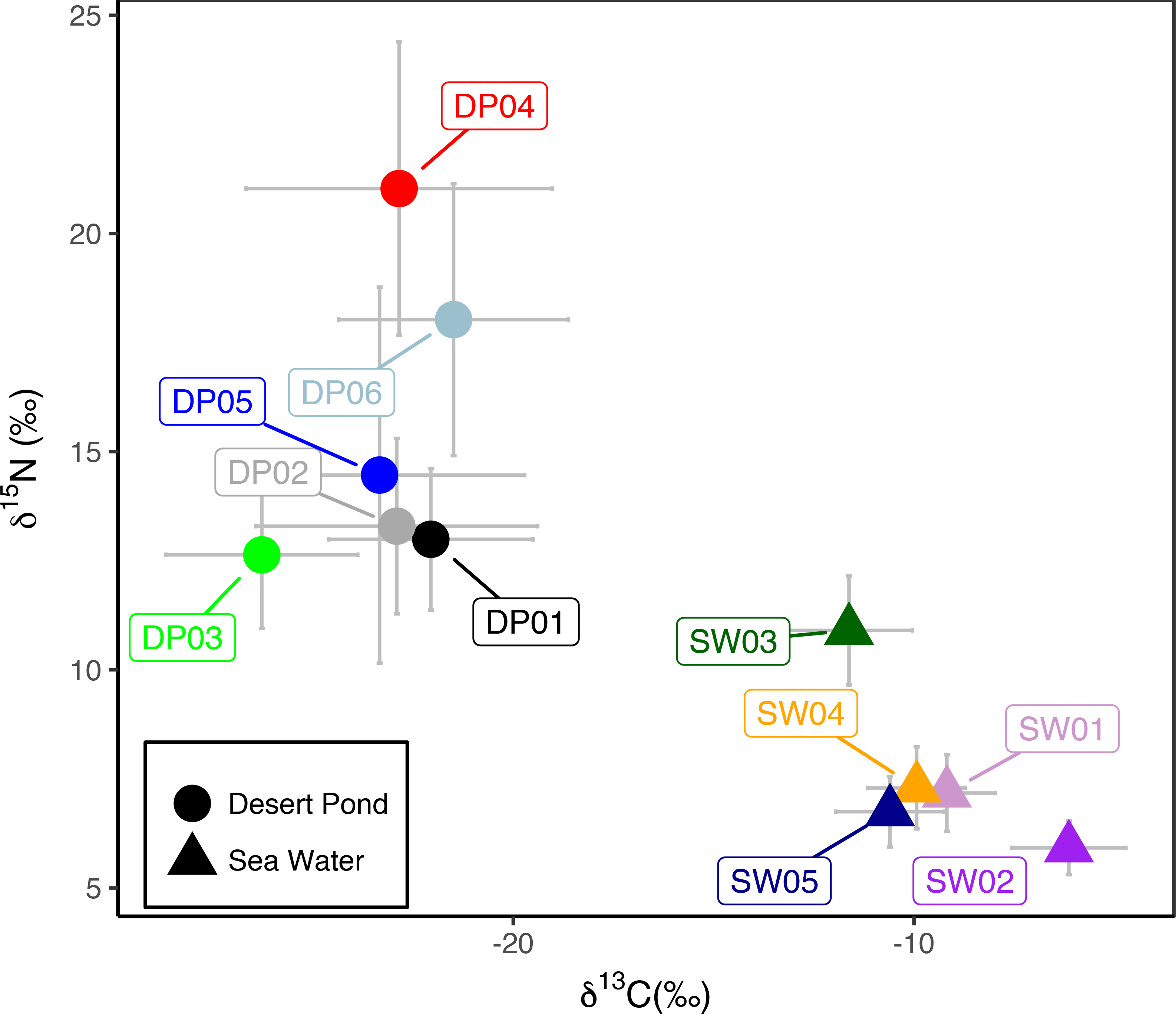
Scatterplot of principal component analysis evaluating the genetic structure between all analyzed *Aphanius dispar* individuals based on 5,955 loci. Eigenvalues represent the amount of genetic diversity shown by each principal component. a) PCA of principal components 1 and 2. b) PCA of principal components 1 and 3.

Since annual rainfall volume tends to increase from the north towards the south (i.e. heavier and more frequent rainfalls towards the south (El Kenawy & McCabe, 2016)), the likelihood of genetic connectivity might also be expected to increase amongst the southern sample sites. In fact, with the exception of DP4, a decrease in genetic distance between locations from the northern area to the south one was found. Moreover, in the southern parts of the study area, less genetic distance was found between desert pond and the nearest seawater location (DP3 with DP4; DP4 with SW4 as well as DP5, DP6 and SW5). This is in accordance with the hydrological map, where water flow is possible from the desert pond sites to the respective wadi (ephemeral river systems appearing during intense rain periods) outlets in the Red Sea. It is not unusual for significant flow events or even flash-floods to occur on an annual, or multi-annual basis (Deng et al., 2015), potentially providing the mechanism behind the observed genetic connectivity within the population clusters. When evaluating the third principal component, large genetic distance between DP4 and most other locations becomes apparent (Figure 5b).

### Putative environmental selection

Possible selective processes to different environmental condition were evaluated using an F_ST_ outlier approach. All desert pond locations were grouped together and compared with all seawater locations, resulting in five outlier loci (Table 3). Based on the population genetic analyses, subsets of desert ponds and genetically close seawater locations were evaluated separately as well. In the northern part, the analysis included two desert ponds (Desert Pond (DP)1 & DP2) and two seawater locations together (Sea Water (SW) 1 & SW2), and resulted in one outlier loci which was also detected during an analysis that included all sites. The comparison of DP4 and SW4 resulted also in one outlier. Comparing DP3 and SW3 recovered one locus only, whereas the three southernmost locations exhibited ten outliers. There is no common overlapping outlier loci detected among the independent salt versus desert pond comparisons. Out of the 21 unique outliers detected in the different subsets of data, 10 could be successfully identified by homology using BLAST, and include several genes involved in immune response and ion channels. The list of NCBI description and accession number can be found in Table 3.

**Table 3:**
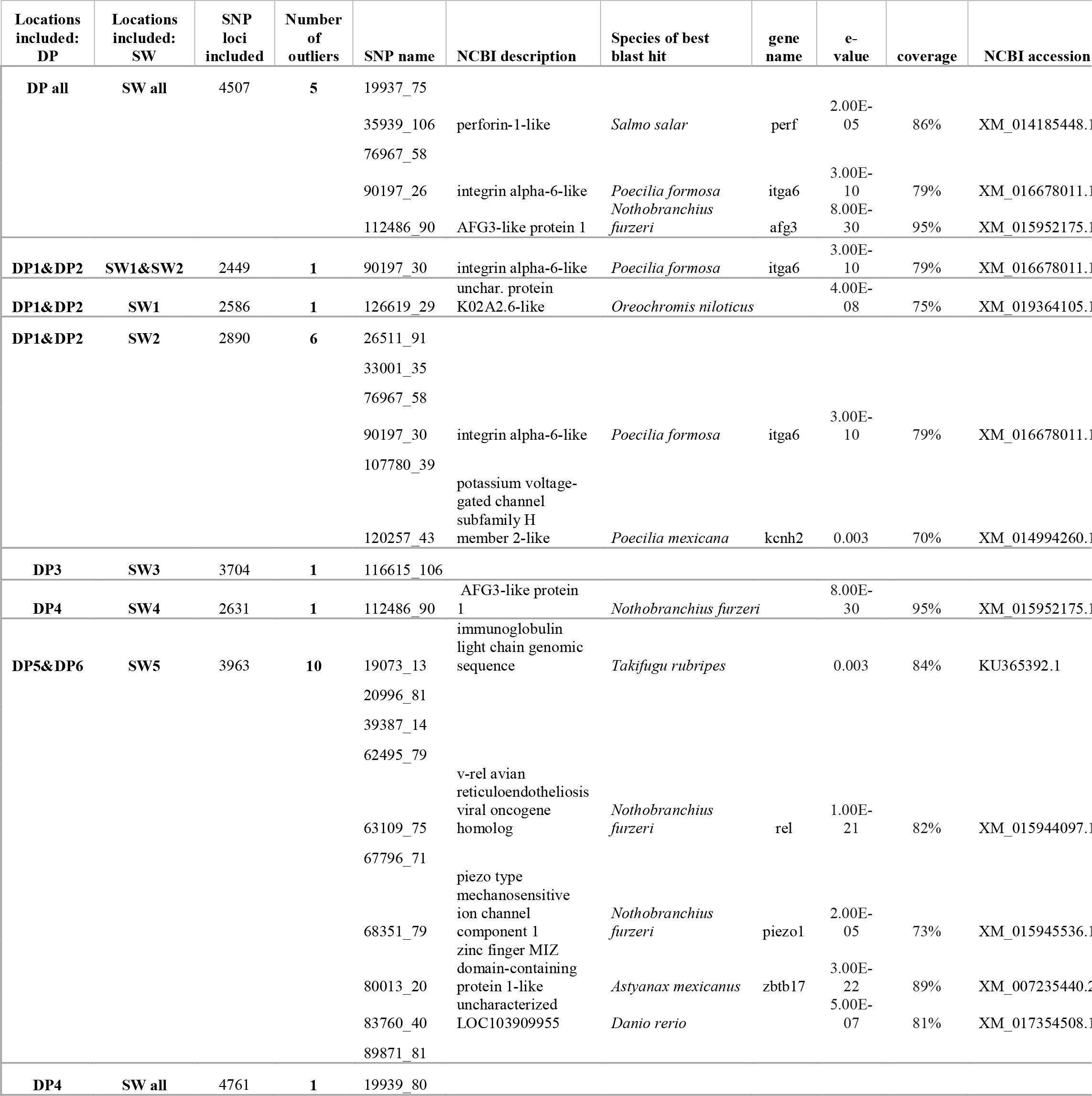
Putative outlier loci between desert pond and red sea locations. Several pairwise comparisons were performed, and the compared locations are indicated under “Locations included”. Sequences were blasted again NCBI and only successful blast hits are listed.

### Muscle tissue stable isotopes

To investigate further into differences between locations and environmental influence on *A. dispar* we analyzed nitrogen and carbon stable isotopes in fish white muscle tissues. These analyses revealed a clear separation between pupfish from inland ponds and saltwater Red Sea habitats (Figure 6). The mean values for δ^13^C and δ^15^N for desert pond fish (DP) ranged between −21.49‰ and −26.28‰ (SD: 2.39‰ to 3.82‰) and 12.63‰ and 21.03‰ (SD: 1.62‰ to 4.31‰), respectively. Saltwater fish (SW) isotope mean values ranged between −6.13‰ and −11.62‰ for δ^13^C (SD: 1.21‰ to 1.58‰) and between 5.92‰ and 10.90‰ for δ^15^N (SD: 0.61‰ to 1.25‰). Overall, DP sites were more depleted in δ^13^C than SW, but had the highest values of δ^15^N. The isotopic signatures of δ^13^C and δ^15^N correlated significantly (p < 0.0001, ρ = −0.85). While gender did not result in a significant covariate (p > 0.05), fish length was significantly correlated with both δ^13^C (p < 0.0001, ρ = −0.39) and δ^15^N (p = 0.0002, ρ = 0.32) values. Our data did not pass the normality test (Shapiro-Wilk test, p < 0.0001) nor the homoscedasticity test (Fligner-Killeen test, δ^13^C p = 0.0003; δ^15^N p < 0.0001). For these reasons, to test the differences between DP and SW isotope contents correcting for fish size, a nonparametric ANCOVA was used. There was a strong water type effect on both δ^13^C and δ^15^N fish content (p < 0.0001). When testing the differences within clusters (cluster 1: δ^13^C p = 0.0037, δ^15^N p = 0.0039; cluster 2: δ^13^C p = 0.0385, δ^15^N p = 0.0352; cluster 3: δ^13^C p = 0.0370; δ^15^N p = 0.0350; cluster 4: δ^13^C p = 0.0121; δ^15^N p = 0.0141), the same result was obtained, with a strong segregation between saltwater and desert pond samples, irrespective of geographic proximity. When comparing between desert ponds, DP3 is significantly different from all other ponds for δ^13^C (p = 0.0064) and DP4 has a significantly larger δ^15^N than the other ponds (p=0.0239). Muscle tissue isotopes analysis therefore suggests that fish in the desert pond have different diets and therefore different ecological niches relative to the populations from the Red Sea coast and some desert ponds have distinct isotope signatures in comparison to other desert ponds.

**Figure 6:**
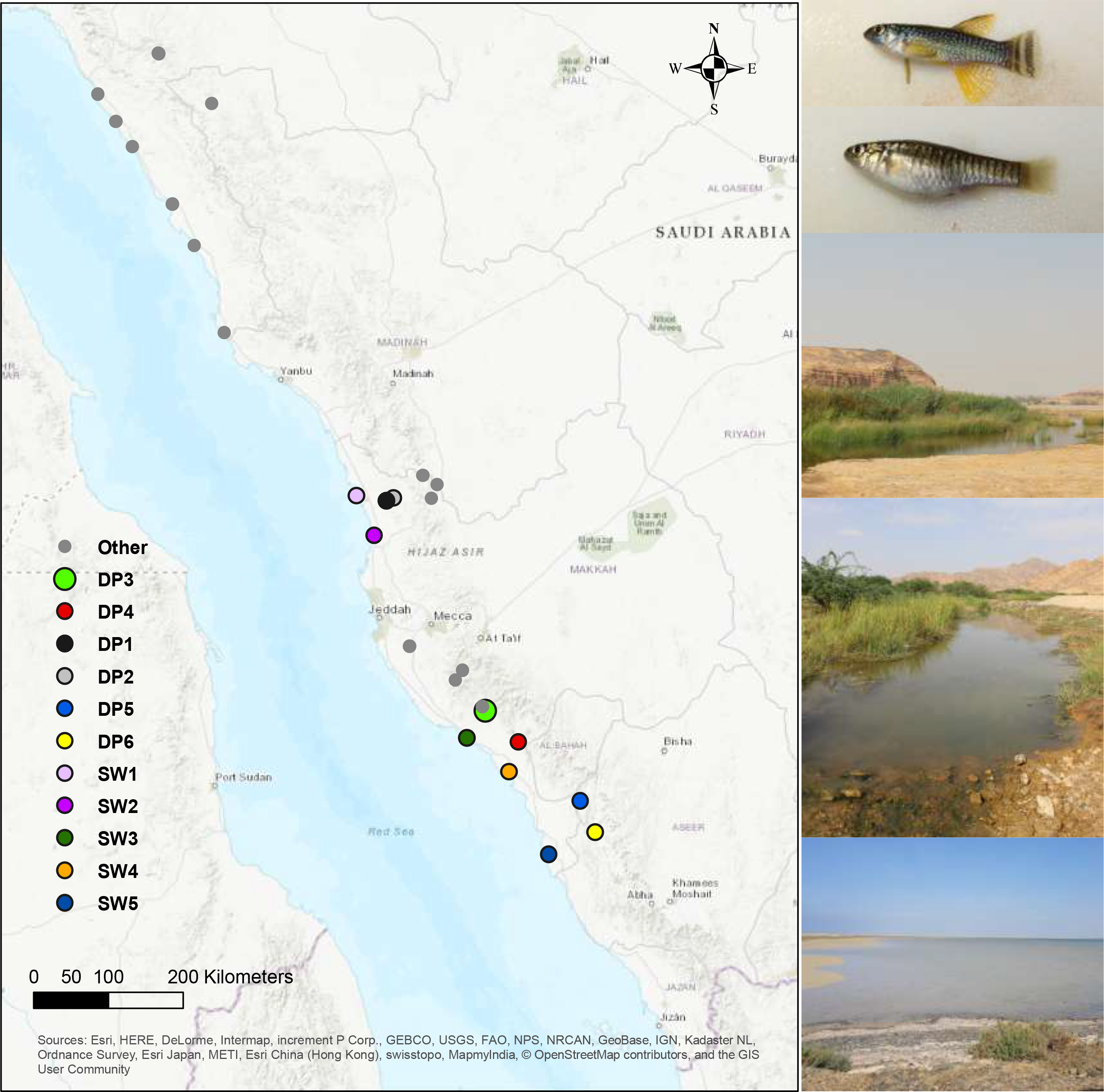
Ratio of stable isotopes of carbon and nitrogen in muscle tissue of *Aphanius dispar*, including fish from brackish water ponds and saltwater in the Red Sea. Error bars represent the standard error per location

## Discussion

Our sampling of *Aphanius dispar* along the Red Sea coastline of the Arabian Desert provides insight to previously uncharacterized sites hosting populations in both near-freshwater water ponds as well as in sheltered coastal marine habitats. In contrast to the often ephemeral habitats occupied by the African turquoise killifish, *Nothobranchius furzeri* (Furness, 2016), the survival and persistence of the Arabian pupfish seems to depend on constant groundwater discharge and irregular rainfall events. Hydrological mapping and genetic analyses reveal that due to sporadic flash floods occurring in the region (Deng et al., 2015) fish may literally be washed from desert ponds out to the Red Sea. The genetic population units comprised an admixture of pupfish from the two different environments, desert pond and seawater individuals. We detected four main genetic clusters, which mostly grouped desert fish with Red Sea individuals based on a latitudinal gradient. Due to little hydrological connectivity among desert ponds, pupfish from the desert (or ‘oases’) are alternately rapidly carried out to very different environmental conditions in the Red Sea along wadis during flash flood events. The change in environmental condition is though not accompanied by population structure, contrary to many other study systems across different environmental gradients with populations differentiating with distance or environment (Sexton et al., 2014).

### Ecological acclimation

Surviving a ‘washout’ event from a desert pond with less than 1 ppt in salinity to the highly saline Red Sea (with an average salinity of 43 ppt) requires a considerable acclimation capacity. Many killifish species are euryhaline and can tolerate large salinity changes in the environment. *Fundulus heteroclitus* has been shown to undergo a large range of physiological changes, from drinking rates to changes in acid base regulation (Wood & Marshall, 1994). Nonetheless, *Fundulus* species, albeit tolerant to large salinity ranges, have diverged into species adapted to specific environmental conditions (Whitehead, 2010). Interestingly, we found the same genetic populations of *A. dispar* living in both desert ponds and seawater. The hypothesized saltwater evolutionary origin of the *Aphanius* genus^19^ provides a potential explanation for the tolerance of *A. dispar* when moved from near-freshwater ponds to a highly saline environment. However, it is not just temperature and salinity that differ between the desert ponds and the Red Sea coastal environments. We also see an ecological divergence in tissue stable isotopes between desert and seawater sites, even from within the same genetic population unit. The seawater fish muscle tissues were depleted in ^15^N and more enriched in ^13^C than their freshwater counterparts. Similar signatures have previously been found when comparing freshwater and saltwater fish, although the fish species in each of the environments differed (Fuller et al., 2012; Robson et al., 2016). Similarly, for anadromous species such as salmon that travel from fresh to saltwater, an increase in ^13^C is exhibited when resident in seawater habitats (Litz et al., 2017). Higher levels of ^15^N, found here for the desert ponds, is often correlated with higher trophic hierarchy levels (Zanden & Rasmussen, 1999). For *A. dispar*, however, it is difficult to isolate the potential influences of habitat-related variability in the basal isotopic values of the food web from the potential trophic shifts (McMahon, Hamady, & Thorrold, 2013; Post, 2002).

### Genetic divergence

Environmental differences leading to ecological divergence can drive adaptation and eventually speciation (Arnegard et al., 2014). Despite large ecological difference in prey and habitat niche, there is little genetic difference between the desert pond fish and the geographically-corresponding seawater fish. This could give a hint that ‘washout events’ occur frequently enough to create enough migration from the desert ponds to the seawater sites, or that, as previously shown in other killifish species (Whitehead, Roach, Zhang, & Galvez, 2012), the Arabian killifish has a large capacity for rapid and long-lasting plastic responses to environmental change. We found only five putative outlier single nucleotide loci that could be under selection between all desert pond inhabitants and seawater fish, and only three that can be functionally annotated. Perforin 1 (*prf1*), a gene to which one of the outliers was successfully blasted (see Table 3), has key functions in the immune response and forms part of killer T-cells, indicating a potential adaptive change to the highly saline environment in the Red Sea. This particular function has been seen to be conserved in many fish species (Nakanishi, Toda, Shibasaki, & Somamoto, 2011; Toda, Araki, Moritomo, & Nakanishi, 2011). Interestingly, in wild salmon, perforin-mediated apoptotic processes were important in survival when migrating back to freshwater spawning grounds from seawater (Miller et al., 2011). Environmental changes are also associated with a SNP in the ATPase Family gene 3 (*afg3*) previously attributed to stress and biosynthesis and with an increased expression in trout after a starvation period (Rescan et al., 2007). A hint towards its importance in seawater acclimation could be related to an increase in mitochondrion-rich cells and ATPase activity due to a physiological demand of acid-base and ion regulation in saltwater (Lee, Hwang, Shieh, & Lin, 2000).

One of the desert ponds, the Al Lith hot spring (site DP3), had high temperatures (>38°C) and a different chemical signature, such as a high amount of silica. Life in such hot water could potentially provoke adaptive signals in the genome. However, only one putative outlier was found for the inhabitants of the hot spring in comparison to other desert ponds. therefore hinting to phenotypic plasticity, as for the case of the Magadi tilapia (*Alcolapia grahami*). *Alcolapia grahami* lives in hot springs in Kenya and was recently described to have the highest upper critical temperature recorded for a fish (45.6 ^°^C (Wood et al., 2016)). Despite the extreme environmental conditions, genetic studies did not find population differences when compared to tilapia of less extreme environmental conditions at close proximity (Wilson et al., 2004; Zaccara et al., 2014). Low numbers of putative outlier SNPs were also detected, suggesting some ongoing gene flow and admixture (Ford et al., 2015), which could be the case for our *A. dispar* samples from the Al Lith hot spring. Despite the lack of loci under selection, connectivity between desert ponds is low and there is limited gene flow, as indicated by large genetic distances between the hot spring and other desert ponds, suggesting isolation and divergence between these habitats. For the pupfish in the hot spring, we could detect a differentiated ecological signature in tissue stable isotopes compared to the other desert ponds. Although contradicting the findings on the Magadi tilapia, where the hot spring site revealed carbon isotope tissue enrichment (Kavembe, Kautt, Machado-Schiaffino, & Meyer, 2016), *A. dispar* from the Al Lith hot spring have lower values of ^13^C in their muscle tissue. One possible explanation for this result might be the turbidity of the hot spring site, due to a large amount of suspended silica, as carbon stable isotope depletion was previously associated with turbidity in the environment (Nahon et al., 2013).

### Anthropogenic impacts on desert fish populations

Besides natural environmental conditions that are reflected in the isotopic signature of the pupfish, anthropogenic impacts can also be detected. In the case of DP4, we found an isotopic as well as genetic signature of human disturbance. Fish tissues here are more ^15^N enriched in comparison to other locations, which indicates an accumulation of the heavier isotope element possibly due to isolation of this pond (Amundson et al., 2003; Szpak, 2014). In this site, there is evidence of agricultural activity, most likely utilizing groundwater in the area, which in turn might lower the water table and disconnect this location from others. Furthermore, there is a dam structure 15 km upstream of DP4 most likely restricting the water availability downstream. Even if topographic mapping shows a hydrological connection between DP4 and SW4, it seems that human interference here inhibits any flow of water to the sea. This hypothesis would explain the genetic differentiation for this particular site, which seems to be undergoing a population bottleneck. The fish in this site have in fact an increased number of private alleles and low genetic diversity, indicated by low heterozygosity. Even though the inbreeding coefficient was is low, this result is most likely due to low genetic differentiation within the population used in the calculation. Pairwise kinship reveals most individuals within this site to be highly related, and the effective population size is very low. It therefore seems that use of water in this area has isolated this population, restricting its gene flow and its resilience.

In the northern part of the Saudi Arabian coast, *A. dispar* in desert ponds and saltwater locations do not cluster together genetically, as it is seen in the southern parts, albeit hydrological connectivity potential. Here the desert pond fish are genetically closer to each other, as are the seawater fish, with a clear division between desert and Red Sea sample sites. There are two plausible reasons for this disconnection. The northern regions receive less rainfall (El Kenawy & McCabe, 2016) and hence any hydrological connectivity between the desert and the sea will be much lower. The observed strong genetic divergence though is most likely caused by the ‘upstream’ construction of the Rabigh Dam (completed 2008), which was built for municipal water supply and flood control. Hence, even with rain events the water, and therefore the pupfish, can no longer reach the sea. For this reason, *A. dispar* populations are now diverging without the presence of gene flow through new migrants. A similar anthropogenic impact was found for desert fish of the Colorado River area, where natural flooding occurred regularly until the construction of dams that drastically changed the water availability and had a major impact on the distribution of desert fish (Hillyard, Podrabsky, & van Breukelen, 2015).

Fish living in desert regions have long been a conservation concern, with a large number of such species being under threat or endangered, often due to the expansion of desert agriculture and increasing global temperatures (Van Haverbeke et al., 2013). Although the Dead Sea subspecies *(A. dispar richardsoni)* is considered endangered (Goren, 2014), *A. dispar* itself is not considered to be endangered because it has stable populations widely distributed throughout the Arabian region (Freyhof, 2014). The species’ capacity to acclimate and survive challenging environmental fluctuations likely plays a major role in its success in this region. However, our results show that despite the large capacity of *A. dispar* to acclimate and adapt to different environments and defy the constraints of living in restricted desert environments, anthropogenic water use can dramatically alter the population dynamics of the Arabian pupfish.

## Acknowledgments

This study was supported by the King Abdullah University of Science and Technology (KAUST). We are very grateful to the KAUST Coastal and Marine Resources Core Lab and KAUST Government Affairs for their support in finding desert ponds, obtaining permits and aiding in the field. We thank the KAUST Integrative Systems Biology Laboratory and the KAUST Biosciences Core Laboratory for support and assistance.

## Author contributions

C.S. designed and managed the field collection. C.S, L.C.B & J.N. performed the sample collection. L.C.B. and J.N. with the supervision of C.S. prepared the samples for DNA sequencing and tissue isotope measurements. M.L.B. provided reagents. J.N. performed chemical water analyses and L.C.B. analyzed the tissue isotope data. S.D.P., Y.A.L. and M.F.M. created hydrological mapping and analyzed water isotopes. C.S. analyzed the sequencing data, performed population genetic analyses with help from J.S. and integrated all datasets. C.S. & T.R. wrote the paper; all authors provided input to and approved the final version of the manuscript.

## Additional information

Raw sequencing data have been deposited on NCBI under BioProject ID PRJNA311159. Final SNP vcf file is available as Supplementary Material. Correspondence and requests for materials should be addressed to T.R. or C.S.

