## Supplementary Figures for "Desert fish populations tolerate extreme salinity change to overcome hydrological constraints"

### Washed out to sea: Desert fish populations tolerate extreme conditions to overcome hydrological constraints

#### Supplementary Materials

Supplementary Figure 1

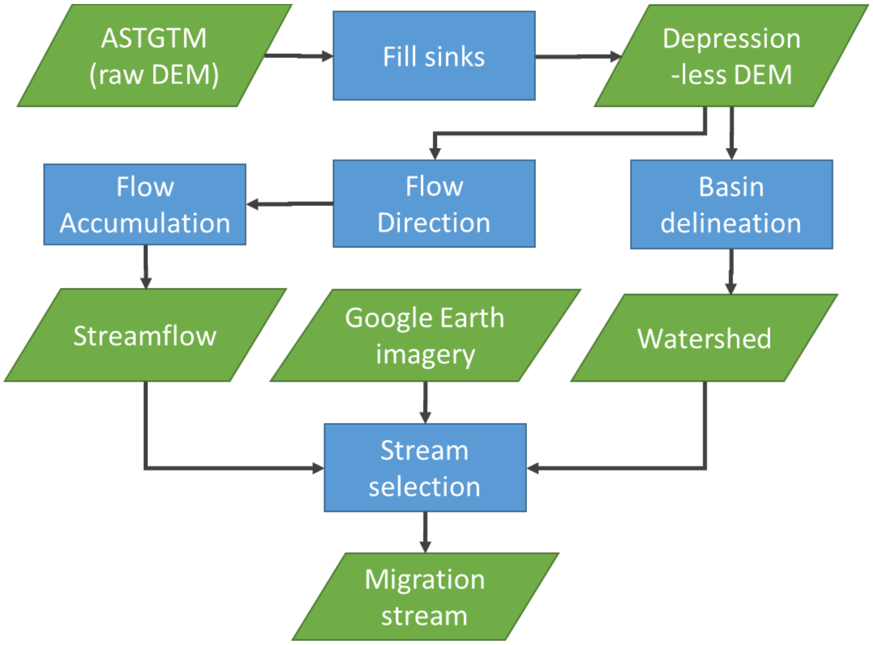


Supplementary Figure 1: Flow process chart of delineating watershed boundaries and stream networks, from the digital elevation model (DEM).

*Water isotopes analysis*

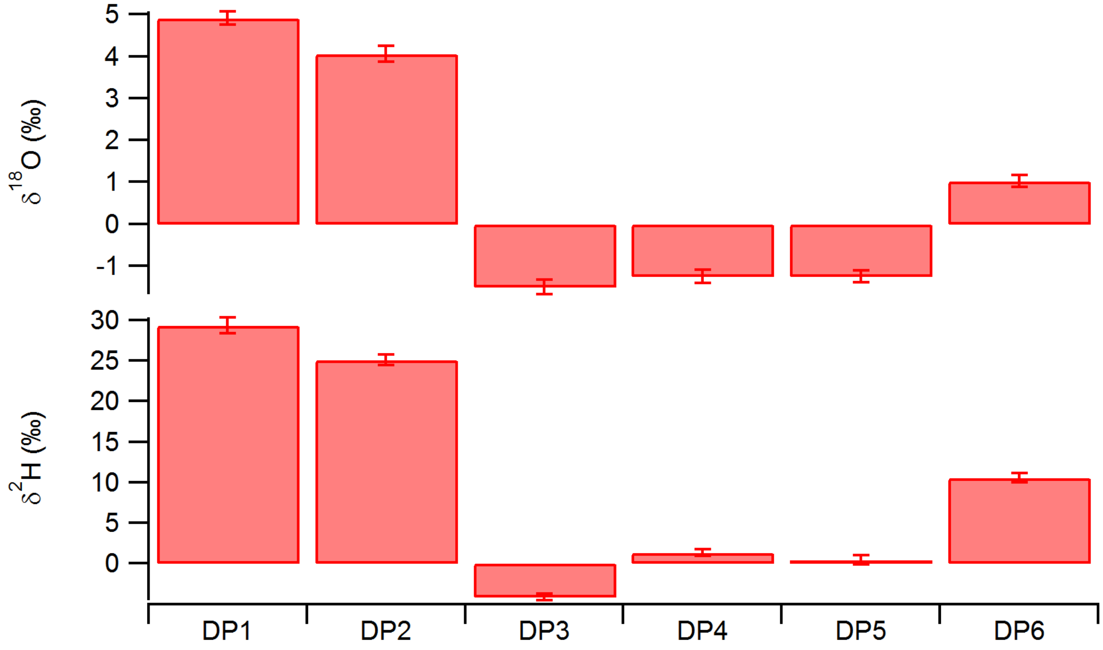


Supplementary Figure 2: Summary of isotope measurements of each sample

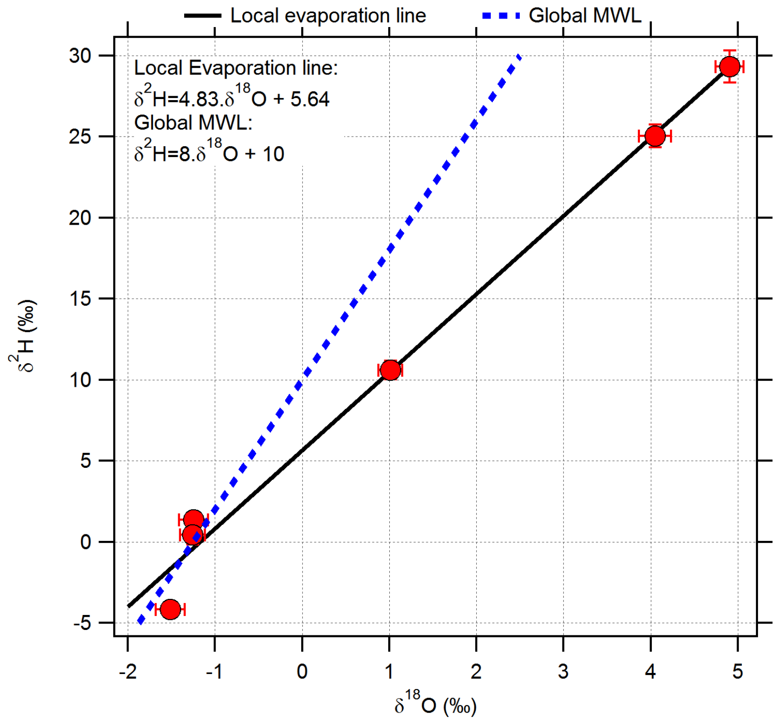


Supplementary Figure 3: The relationship between $\delta^2H$ and $\delta^{18}O$. Note the slope is much less than 8 indicating evaporative processes.

*RAD sequencing processing and SNP marker selection*

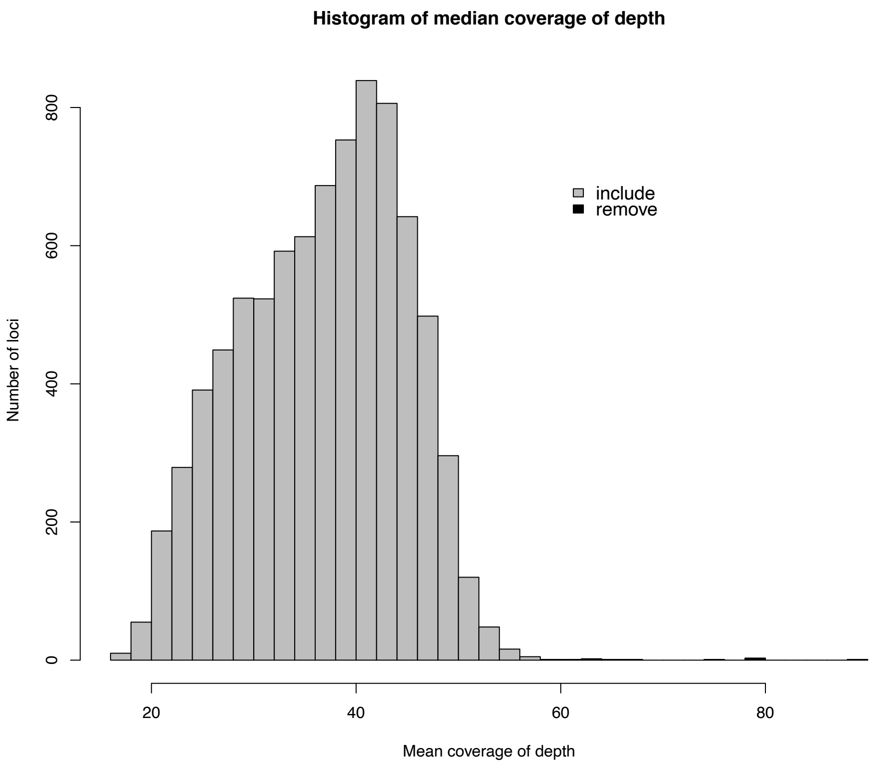


Supplementary Figure 4: Distribution of sequence coverage depth for each loci after filtering with *populations*. Most of the loci were included in the subsequent analyses, some had an increased depth of coverage, possibly hinting to paralog loci and were therefore removed.

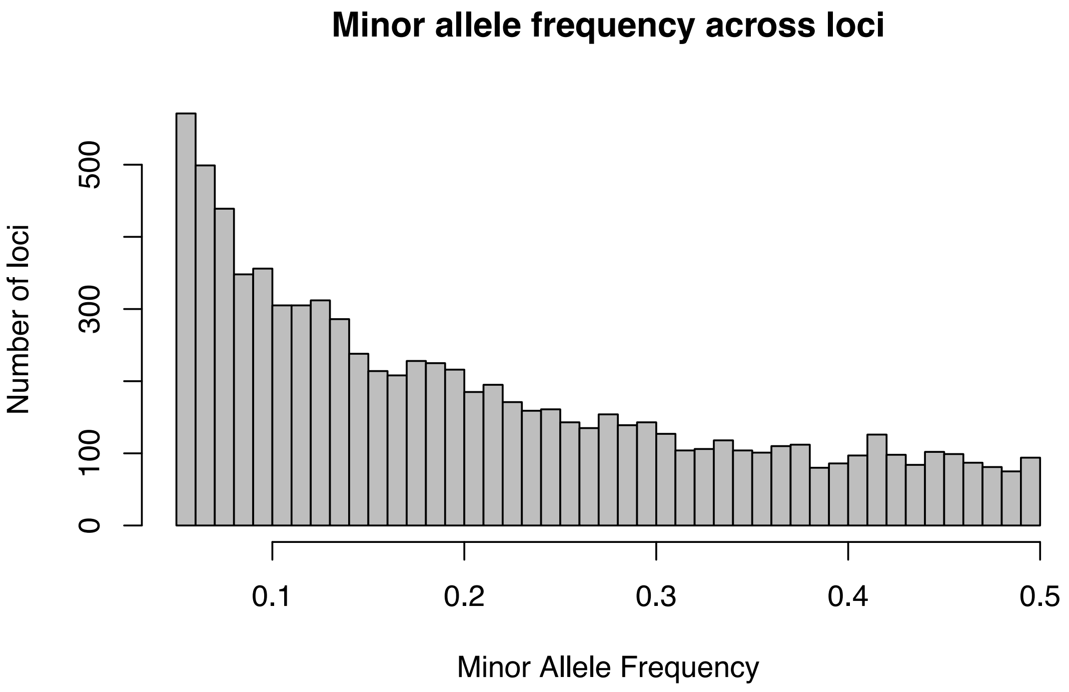


Supplementary Figure 5: Distribution of minor allele frequencies across all selected loci.

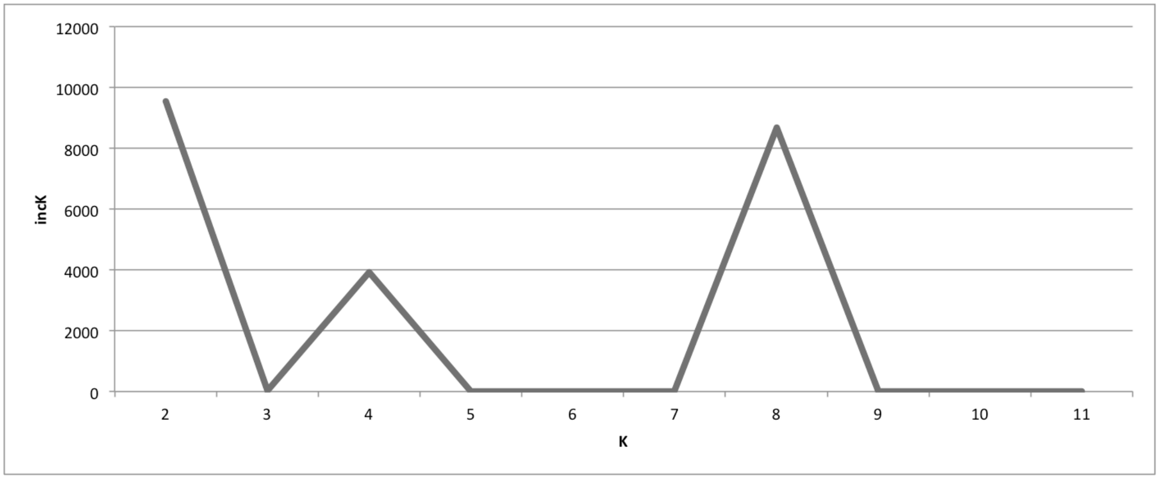


Supplementary Figure 6: Post-hoc test results (Evanno et al.) for the K-cluster in the STRUCTURE analysis. X axes= number of K. Y axes= increment of K
